# The Evolutionary Dynamics of Hyperparasites

**DOI:** 10.1101/2021.12.01.470853

**Authors:** Graham R Northrup, Steven R Parratt, Carly Rozins, Anna-Liisa Laine, Mike Boots

## Abstract

Evolutionary theory has typically focused on pairwise interactions, such as those between hosts and parasites, with relatively little work on more complex interactions including hyperparasites: parasites of parasites. Hyperparasites are common in nature, with the chestnut blight fungus virus CHV-1 a well-known natural example, but also notably include the phages of important human bacterial diseases. Theory on hyperparasitism has mostly focused on their impact on the evolution of virulence of their parasite host and relatively little is known about evolutionary trajectories of hyperparasites themselves. Our general modeling framework highlights the central role the that ability of a hyperparasite to be transmitted with its parasite plays in their evolution. Hyperparasites which transmit with their parasite hosts (hitchhike) will be selected for lower virulence, trending towards hypermutualism or hypercommensalism and select against causing a reduction in parasite virulence (hypovirulence). We examine the impact on the evolution of hyperparasite systems a of a wide range of host and parasite traits showing, for example, that high parasite virulence selects for higher hyperparasite virulence feeding back into selection for hypovirulence in the parasite. Our results have implications for hyperparasite research, both as biocontrol agents and for understanding of how hyperparasites shape community ecology and evolution.

## Introduction

Evolutionary theory is well developed for simple pairwise examples of evolution such as predator/prey (Abrams 2000), host/mutualist (Akçay 2015) and in particular host/parasite (Best et al. 2009) interactions, but only rarely considers interactions within communities of interacting species (Thompson 1999, Brodie and Ridenhour 2003). The coevolution of hosts and pathogens is well studied both theoretically (Sasaki 2000, Anderson and May 1983, Best et al. 2009, Boots et al 2014) and empirically (Brockhurst and Koskella 2013, Ebert 2008, Laine 2006, Gandon et al. 2008, Gómez and Buckling 2011), providing a solid framework for understanding of the key evolutionary drivers of their life history such as virulence, resistance and recovery (Cressler et al 2016). However, far less is known about how host-parasite interaction traits evolve when we account for the biotic community they are embedded in (Alizon and Van Baalen 2008, Osnas and Dobson 2012, Hall et al. 2020). There are clearly analytical and conceptual challenges to moving beyond pairwise interactions, but one important but tractable multi-parasite interaction is hyperparasitism (Parratt and Laine 2016). Hyperparasites are parasites that infect hosts which are themselves a parasite of another host, they are widespread, and diverse in nature (Parratt and Laine 2016) and as such they represent an important but understudied three-way interaction. In this paper, we develop a general evolutionary framework for hyperparasites referring throughout to the three players in this system as the host, the parasite, and the hyperparasite.

Modeling of hyperparasite systems is motivated not only by our general interest in understanding evolution beyond pairwise interactions, but also due to the number of diverse and impactful hyperparasitic systems in nature (Parratt and Laine 2018, Choi and Nuss 1992, Vandermeer et al. 2009). Hyperparasites exist across a diversity of systems and types of hosts in nature. A well-known example is the chestnut blight fungus, *Cryphonectria parasitica*, a parasite of chestnut trees that also plays host to the hyperparasitic CHV-1 mycovirus which has been shown to reduce the virulence in chestnut trees caused by *C. parasitica* (Choi and Nuss 1992). There are fungal hyperparasites such as *Ampelomyces quisqualis* which parasitizes powdery mildews (Kiss et al. 2004) and of course a wide range of bacteriophage hyperparasites of bacterial pathogens (Cruz-Flores et al. 2016). In addition, there is considerable interest in the application of hyperparasites in biocontrol. In particular, phage therapy has already been used to treat antibiotic resistant (AMR) infections in several cases, including in particular in cystic fibrosis patients (Kortwright et al. 2019), and the hyperparasitic mycovirus has been successfully used to control chestnut blight (Rigling and Prospero 2017). More generally, it is now thought that the diversity of hyperparasites may be an important component of observed pathogen virulence patterns (Parratt and Laine 2018). Understanding the ecological and evolutionary dynamics of these systems is therefore important both due to their role in natural systems and their potential use in therapeutics (Wang et al. 2020).

Current hyperparasite theory has often been focused on specific systems and therefore makes system specific assumptions about the impact of the hyperparasite on its host (Taylor et al. 1998, Morozov et al. 2007) although Sandhu et al. 2021 has recently pursued more general results. Here we focus on the unique characteristics of a hyperparasite; a hyperparasite decreases the fitness of its own host (a parasite), a hyperparasite has the potential to impact the effect of the parasite on its host, including inducing hypo or hyper parasite virulence, and a hyperparasite may increase the death rate of the parasite (hyperparasite virulence), essentially clearing the parasitic infection and leading to an uninfected host. Understanding the conditions under which a hyperparasite may evolve to induce hypovirulence in their parasite host is of particular interest, as this will elucidate the effects of hyperparasites in nature or when they are introduced as biocontrol agents. Importantly, we examine the impact of the hyperparasite’s ability to be transmitted along with the parasite when it infects a new host (hitchhiking).

Holt and Hochberg (1998) proposed a hyperparasite model with a focus on how this important tritrophic interaction impacts the ecological stability of communities. The model assumed the same death rates in parasitized and hyperparasitized hosts (*c_α_* = 1 in our general framework defined below) and no hyperparasite virulence, meaning that hyperparasitized hosts do not recover to be fully susceptible. There is therefore effectively no damage from hyperparasite infection to the parasite. Their models also assume 0% hitchhiking not allowing direct hyperinfection of a completely susceptible host and their obligate hyperparasite model also does not allow transmission of hyperinfected parasites (*c_β_* =0 in our general framework defined below). The model predicts that hyperparasitism should select for higher parasite virulence, which is equivalent to the prediction of the related co-infection model (May and Nowak 1995). Taylor et al. (1998) focus on the evolution of hypovirulence in a model focused on the *C. parasitica* and CHV-1 system. They assume 100% hitchhiking, meaning that an infection event between a susceptible host and a hyperinfected host will always transmit both pathogens to the new host but assume reduced transmission of a hyperparasitized parasite (*c_β_* < 1 in our general framework) and have a parameter which measures the difference in transmissibility that occurs through hitchhiking (S and H contact, called vertical transmission in Taylor et al. 1998). Recently, Sandhu et al. 2021 examined the coevolution of hyperparasites and parasites with an explicit adaptive dynamical model. They predict that hyperparasites will always select for higher parasite virulence in their host, although they can still act as excellent biocontrol agents as they reduce parasite prevalence, and also show the interesting possibility of evolutionary suicide. This work assumes there is no recovery from hyperparasitized to susceptible hosts as a result of hyperparasites exploiting the parasite and killing their ‘host’ and therefore, they did not examine hyperparasite virulence directly and furthermore, they assumed 100% hitchhiking. As such we lack a general understanding of the evolution of hyperparasites with the existing theory not focused on key traits such as their virulence and assuming very different degrees of hitchhiking.

We build on this previous work, by creating a general framework for the evolution of hyperparasites, incorporating virulence of the hyperparasite on the parasite, and in particular specifically examines how the assumption of hitchhiking affects evolutionary outcomes. While previous general evolutionary theory has concentrated on the impact of the hyperparasite on parasite evolution, we explicitly model the evolution of key hyperparasite traits including its virulence. We emphasize that the degree of hitchhiking is a key characteristic of any particular hyperparasite system with fundamental implications to the evolutionary outcome.

## Methods

In order to understand the evolutionary dynamics and outcomes of hyperparasites, we build a general modelling framework consisting of three ordinary differential equations that capture the infection-status of a host population. The system of equations describes the dynamics of uninfected hosts (S), hosts infected with a parasite (I), and hyperinfected hosts (H). Hyperinfected hosts are those that are infected by the intermediate parasite which is itself infected with the hyperparasite (see Figure 1). Here we consider obligate hyperparasites only, which require the parasite present in the host.

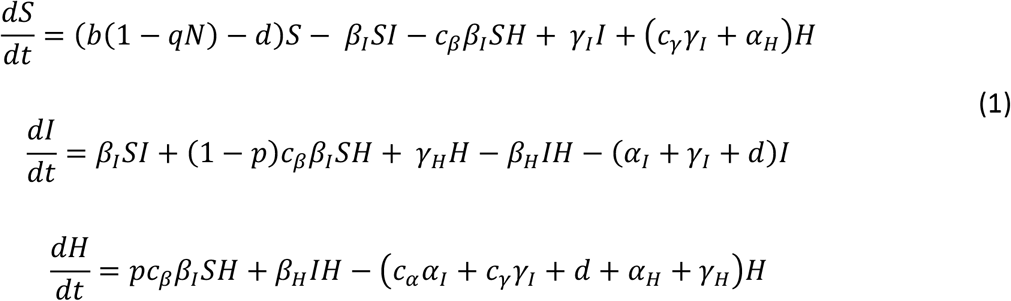

**Figure 1:**
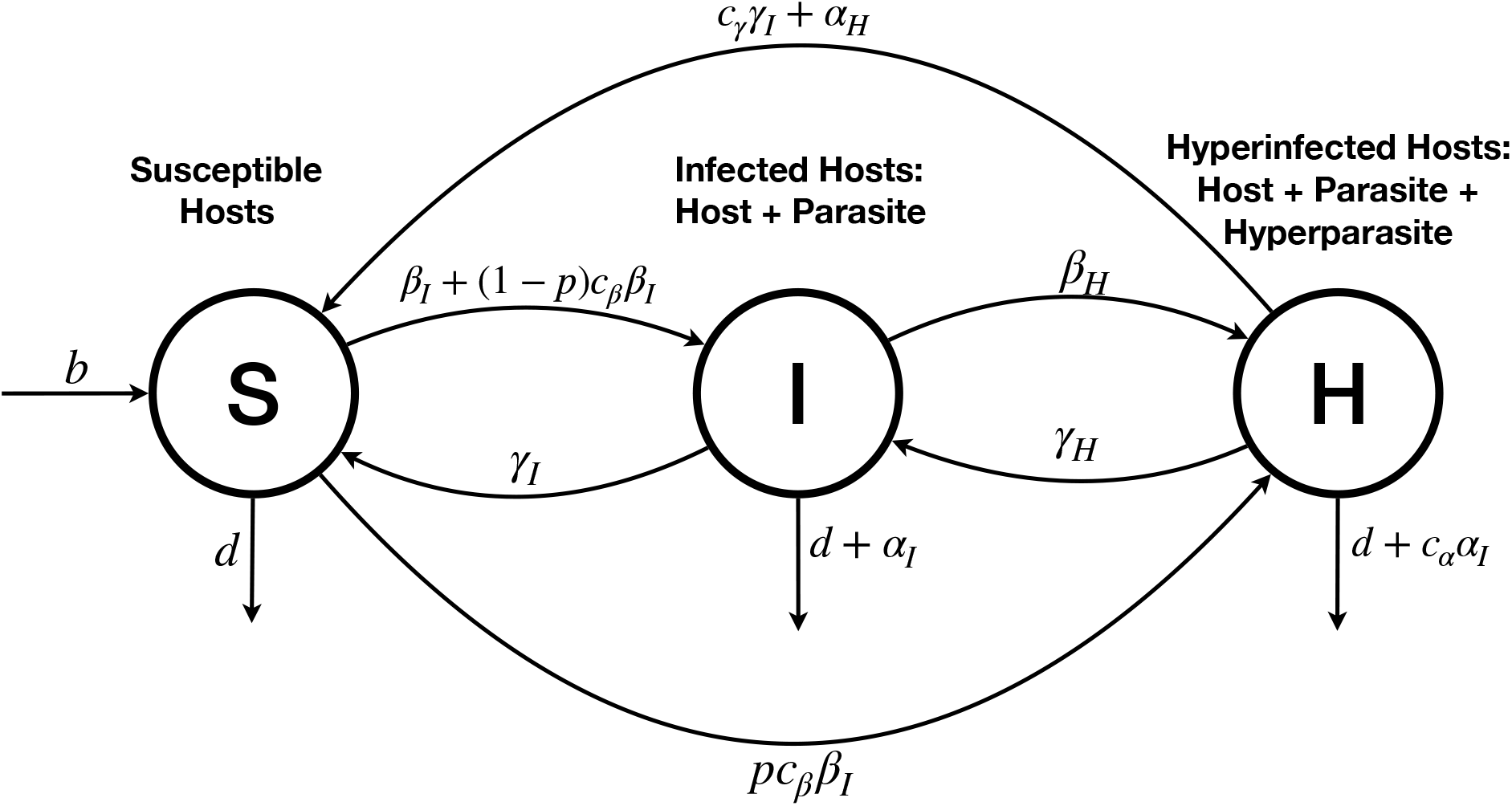
Diagram of model for hyperparasite system. Susceptible hosts in S, singly infected hosts in I, and hyperinfected hosts in H. α_I_ & α_H_ describe the virulence of the parasite and hyperparasite respectively. β_I_ & β_H_ describe the transmission of the parasite and hyperparasite respectively. γ_I_ & γ_H_ describe the recovery rates for the parasite and hyperparasite respectively. The parameter p governs the probability a hyperinfected parasite brings its hyperinfection to a susceptible host (hitchhiking). Parameters of the form c_δ_ (where δ is an parasite natural history trait) reflect the change in the parasite parameter δ_I_ as a result of hyperinfection.

**Figure 2:**
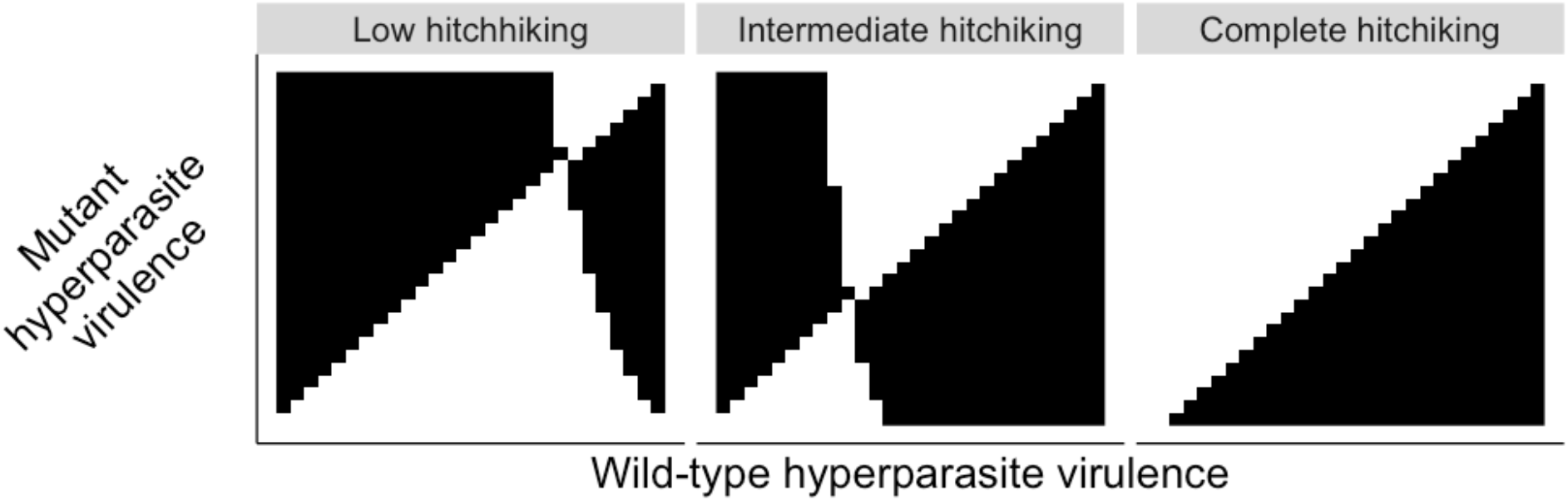
Pairwise invasability plots (PIPs) showing the evolutionarily stable strategy (ESS) for hyperparasite virulence α_H_ under varying values of the parameter p – the propensity to hitchhike -. A black square shows a parameter combination where the mutant hyperparasite will invade, and a white square is one where the wild type hyperparasite will outcompete. This is done under a virulence transmission tradeoff as described in the text. b = 1, d = 0.1, q = 0.0005, α_I_ = 0.05 (parasite virulence), β_I_ = 0.01, γ_I_ = 1, α_H_ = 0.05 (hyperparasite virulence), β_H_ = 0.05, γ_H_ = 0.01, c_α_ = 1, c_β_ = 1, c_γ_ = 1. This is consistent with a hyperparasite that has neutral effects on the life history traits of the parasite.

Susceptible hosts experience density dependent growth, with *b* as the growth rate and *q* as the crowding term. All hosts experience a natural death rate *d*, regardless of infection status. Susceptible hosts can become infected by contact with infected hosts at rate *β_I_*. They can also become infected by hyperinfected hosts as well, leaving the hyperparasite behind during transmission, at rate (1 – *p*)*c_β_β_I_* reflecting the hyperparasite’s ability to affect parasite fitness, *c_β_*, and the probability a susceptible host interacting with a hyper infected host creates an infected host (rather than a hyperinfected host), 1 – *p*. Infected hosts can also become hyperinfected through contact with hyperinfected hosts at rate *β_H_*. Infected and hyperinfected hosts can die due to parasite virulence *α_I_* (again hyperinfected hosts may have a different rate this time according to *c_α_*). Infected hosts recover, returning to the susceptible class, at rate *γ_I_* while hyperinfected hosts recover and return to the infected class at rate *γ_H_* and to the susceptible class at rate *c_γ_γ_I_* (recovering from the parasite also de facto clears the hyperparasite from the host). In this same sense hyperinfected hosts can also ‘recover’ to the susceptible class by virtue of the hyperparasites virulence against the parasite.

An important concept in our model is that hyperparasite virulence *α_H_* takes host individuals from H to S, ‘killing’ the parasite but not the host. This is because if the hyperparasite does in fact kill its ‘host’, the parasite, and therefore functionally clears the base host of its original parasite infection. In this way hyperparasite virulence, *α_H_* increases the recovery rate of the host in addition to natural recovery *γ_I_*, where the host is clearing the parasite. So, while the standard parasite virulence *α_I_* increases the mortality of its host, hyperparasite virulence *α_H_* decreases the mortality of the base host. Hyperparasite virulence *α_H_* takes individuals out of H, thus in an equivalent way to *α_I_* it reduces the duration of the hyperparasite infection, but crucially it doesn’t remove hosts from the population rather by allowing them to recover decreasing their mortality. There are previous models which have not implemented virulence of the hyperparasite in this way (Sandhu 2021), but this is the consistent way to use the term virulence, since the hyperparasite is exploiting the intermediate parasite, rather than the base host as might occur in a superinfection model. It is a key unique feature of hyperparasites.

We also introduce a parameter *p*, to capture the propensity for exposure to hyperinfected hosts to infect the susceptible host with both the parasite and then hyperparasite. When hyperparasitized hosts come into contact with susceptible hosts, a proportion *p* become hyperparasitized (they contract the parasite as well as the hyperparasite) while a proportion 1 – *p* lose the hyperparasite and only contract the parasite. This parameter is important as it can fundamentally change the structure of the interaction and the model and emphasizes that there is an important distinction between hyperparasites that “hitchhike” on primary infections and those that are lost at infection (see table 2 for examples). The degree to which a hyperparasite is able to be transmitted with its parasite is a crucial characteristic of the hyperparasite. With *p*=0 the hyperparasite transmission is decoupled from that of its parasite host, but once there is significant hitchhiking there is a fundamentally different relationship between the fitness of the hyperparasite and parasite.

**Table 1:** Description of model parameters separated by general class of parameter

| PARAMETERS | DESCRIPTION |
| --- | --- |
| $b, d, q$ | Birth rate, natural death rate, crowding term (susceptibility to crowding) of the host |
| $\alpha_I, \beta_I, \gamma_I$ | Virulence (increased host mortality due to infection), transmission, and recovery rates of the parasite |
| $\alpha_H, \beta_H, \gamma_H$ | Virulence, transmission, and recovery rates of hyperparasite |
| $c_\alpha, c_\beta, c_\gamma$ | Hyperparasite effects on $\alpha_I, \beta_I, \gamma_I$ parasite virulence, transmission, and recovery, respectively |
| $p$ | Probability of a S+H contact producing a hyperinfected host (as opposed to an infected host): Hitchhiking |

**Table 2:**
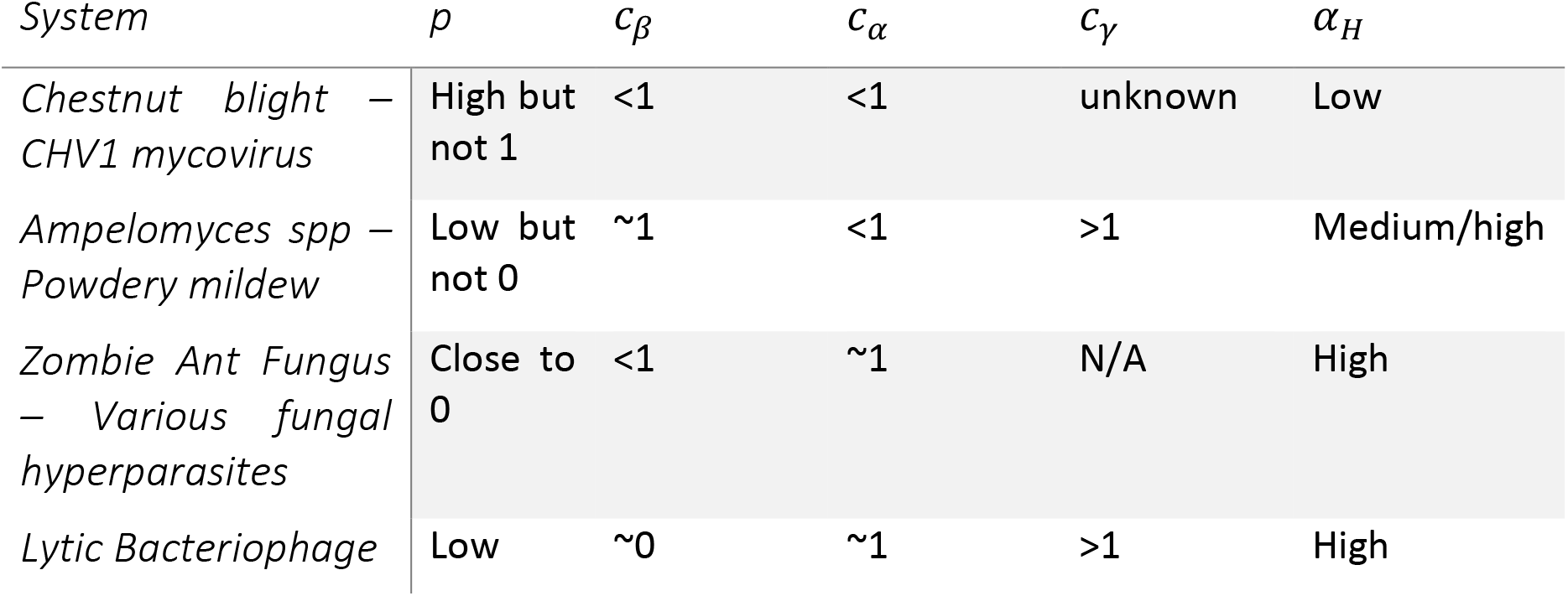
Rows showing estimated system parameters for several parasite-hyperparasite systems, chestnut blight-CHV1 mycovirus (Choi and Nuss 1992) (Rigling and Prospero 2017), Ampelomyces spp-powdery mildew (Tollenaere et al. 2014) (Parratt and Laine 2018), zombie ant fungus-fungal hyperparasites (Andersen et al. 2012), and lytic bacteriophages (Cruz-Flores et al 2016) (Kortwright et al. 2019).

| *System* | $p$ | $c_\beta$ | $c_\alpha$ | $c_\gamma$ | $\alpha_H$ |
| --- | --- | --- | --- | --- | --- |
| *Chestnut blight* – *CHV1 mycovirus* | High but not 1 | <1 | <1 | unknown | Low |
| *Ampelomyces spp* – *Powdery mildew* | Low but not 0 | ~1 | <1 | >1 | Medium/high |
| *Zombie Ant Fungus* – *Various fungal hyperparasites* | Close to 0 | <1 | ~1 | N/A | High |
| *Lytic Bacteriophage* | Low | ~0 | ~1 | >1 | High |

The parameters *c_α_, c_β_, c_γ_* give us considerable flexibility to parameterize this model for a wide variety of host-parasite-hyperparasite systems since they describe the effect of hyperparasites on their parasite hosts key life histories. Different hyperparasites can affect their hosts’ life histories in various ways, and this general modeling approach allows us to incorporate multiple types of effects of hyperparasites on their parasite hosts. These key traits are *c_β_* (parasite transmission modification), *c_α_* (parasite virulence modification), *c_γ_* (parasite recovery modification). Each of these traits reflect the cost to the parasite of being infected with the hyperparasite which has the potential to make them less able to transmit *c_β_* and easier to recover from *c_γ_* and potentially less impactful on their host *c_α_*. One simple way in which these effects can be understood is that the hyperparasite reduces the growth rate of the parasite within its host. Clearly not all hyperparasites will impact each of these traits, but in principle they are all processes where the hyperparasite reduces the parasites fitness. It is important to note that this impact on the fitness of the parasite is the key assumption that defines the hyperparasite – it is a parasite because it reduces the fitness of its host, which is also a parasite. If a hyperparasite did not impact its parasite host fitness, through one or more of these modification terms or virulence *α_H_*, it would be a hypermutualist or hypercommensal.

An important driver of the evolution of antagonist ecological systems is the feedbacks between ecological and evolutionary dynamics (Govaert et al. 2019, Straus 2014, Ashby et al. 2019, Boots et al. 2009). In order to understand the evolutionary implications of our model, in response to ecological feedbacks, we made use of adaptive dynamics to investigate how mutant hyperparasites with a small change in a trait of interest may or may not outcompete the wild type hyperparasite. This is carried out by introducing a fourth equation to our system (1), which models individuals hyperinfected with a novel mutant hyperparasite, comparing various combinations of wild-type and mutant parameter values to determine which becomes the resident strain in each case. The details of the analysis are explained in the supplemental information. Importantly, the model dynamics are highly nonlinear, so we opted for a numerical approach as closed form analytical results proved tough to obtain. This was carried out in a section of the parameter space where we know the model’s residence equilibrium is stable.

We introduce two tradeoff schemes, the first being a standard virulence transmission tradeoff where 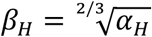. Note that this function is concave down (or saturating) meaning as host (the hyperparasites host in this case i.e. the parasite) exploitation increases, the level of transmission begins to level off as it is also influenced by other factors such as contact rate. We then add to this tradeoff by including the parameters *c_α_, c_β_, c_γ_* as functions of parasite ‘host’ exploitation by the hyperparasite, as well, in line with our biological expectation that these would indeed be affected with *c_α_* and *c_β_* both decreasing towards 0 as host exploitation increased and *c_γ_* increasing as host exploitation increased. In this way we can understand how the nature of a system may create conditions where important effects such as the induction of hypovirulence of the intermediate host are adaptive. We examine a wide range of scenarios to develop a general theory of the evolution of hyperparasites

## Results

We first investigated the existence and nature of the evolutionarily stable strategy (ESS) of virulence. In this analysis we only consider continuously stable points that are the expected endpoint of evolution, for the hyperparasite under the classic virulence transmission tradeoff framework. A key insight is that *p* has a critical impact of the evolutionary stable (ES) virulence, 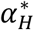, for the hyperparasite and therefore whether the hyperparasite is able to hitchhike with the parasite at infection is critical to the evolutionary outcome. Fundamentally, as the value of *p* decreases, the ES virulence 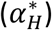 increases. At the maximum value of *p* = 1, when the hyperparasite perfectly hitchhikes, the model selects for avirulence of the hyperparasite or 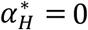. This can be understood because as *p* approaches 1, more arrivals to the hyperinfected class arrive through the *SHβ_I_* transmission term, rather than passing through the infected class first. Conversely as *p* approaches 0, and the hyperparasites stop hitchhiking as frequently, individuals must pass through the infected category to reach the hyperinfected category, thus increasing the relative importance of higher *β_H_* values and thus also higher levels of virulence at the ESS. This result is key to understanding how the fundamental nature of hyperparasite transmission will affect the ESS, as the interplay between *p* and *β_H_* can exert significant control over the system. In particular, hyperparasites will be selected to reduce their own transmission to very low levels when they can hitchhike and become mutualistic with their parasites – leading to hyper commensalism/mutualism.

With the broader tradeoff to include the parameters *c_α_, c_β_, c_γ_* all as functions of host exploitation. Under this more complex tradeoff, we still see that *p* still exerts strong control over the ES virulence of the hyperparasite. Again with *p* = 1 the hyperparasite is selected towards hypermutualism and not impacting the parasite fitness (Fig 3B).

**Figure 3:**
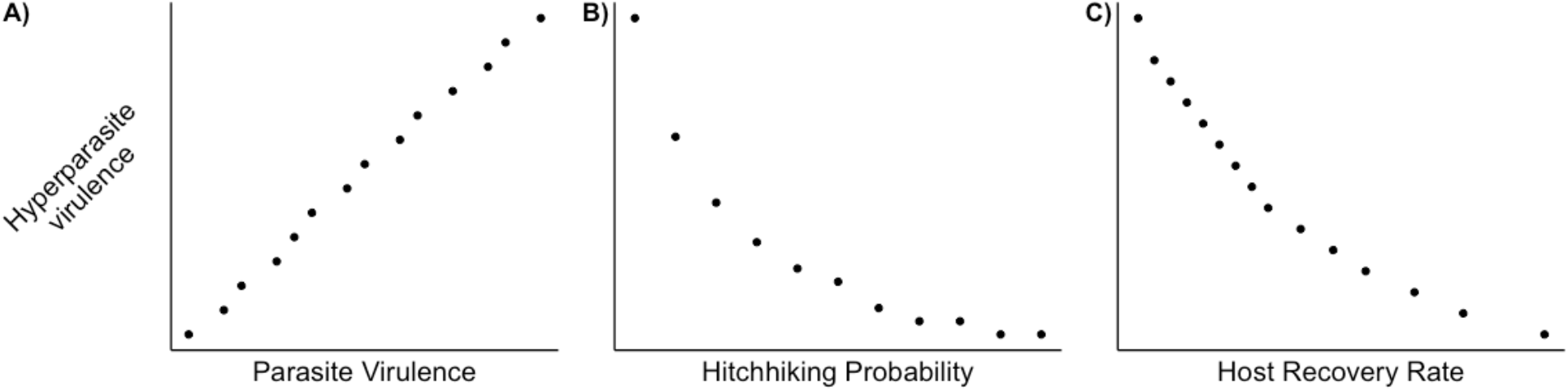
These panels show the relationship between the ES values of 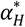 and different values of fixed model parameters. A) is under changing α_I_, B) shows changing values of p, and C) changing values of γ_I_. Non evolving parameter values are otherwise consistent with those used elsewhere.

**Figure 4:**
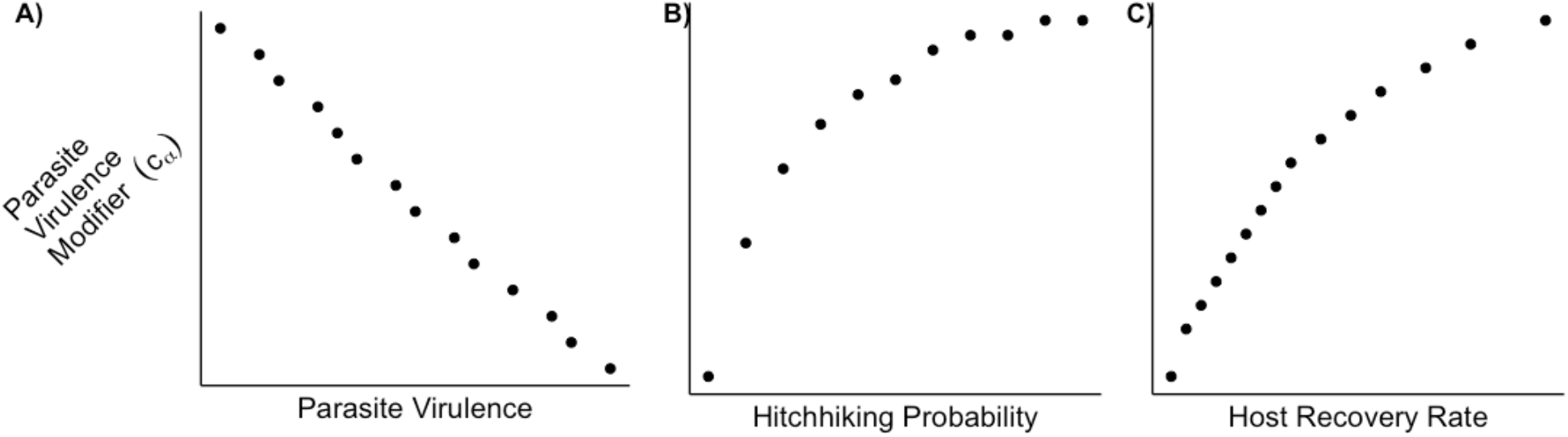
These panels show the relationship between the values of c_α_ associated with the ES virulence and different values of fixed model parameters. A) is under changing α_I_, B) shows changing values of p, and C) changing values of γ_I_. Non evolving parameter values are otherwise consistent with those used elsewhere.

We next examined the importance of the parasite’s life history parameters, to understand how this may influence the hyperparasite life histories including virulence ESS 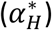. In particular, increasing values of *α_I_* corresponds to an increase in 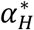. Meaning if the parasite is deadly, its hyperparasite will evolve to become more virulent in turn. Conversely, when increasing the host recovery rate from the parasite, *γ_I_*, this causes a decrease in hyperparasite virulence. These selective forces result in the rate at which individuals leave the H compartment to return to the S class (as both of these are rates of H to S) to be relatively constant, which we hypothesize is some biologically optimal level for the hyperparasite.

These results can also be understood through the lens of *c_α_*, the effect of the hyperparasite on parasite virulence *α_I_*. As shown in Figure 3, subplot A, we can see that increasing the virulence of their host will select for stronger induction of hypovirulence (*c_α_* → 0). This is because the hyperparasite will need to in turn exploit its host more, driving the value of *c_α_* towards 0 as shown. The opposite effect is seen when increasing *p* or *γ_I_*, where *c_α_* values will increase as the ESS shifts accordingly with increased host exploitation in line with the decreasing virulence and transmission previously discussed, according to the hypothesized tradeoff where *c_α_* grows towards 1 as host exploitation decreases to 0.

Finally, we examine how host traits impact the selection on hyperparasite traits. In particular longer-lived hosts selected for less virulent hyperparasites. This is consistent with our understanding of virulence selection as increasing the host death rate would reduce the duration of hyperinfection. This in turn provides selection pressure for more rapid growth and reproduction of the hyperparasite, the source of virulence under our assumptions. This is shown in Figure 5. Furthermore, hyperparasite virulence selection appears unaffected by host crowding over the parameter area searched. Although these types of hosts have higher densities of individuals, facilitating transmission in a density dependent transmission scenario such as our model it appears there is a trade-off between this decreased selection pressure on transmission (which may drive virulence down) and decreased costs of hyperparasite virulence (which may pull it up).

**Figure 5:**
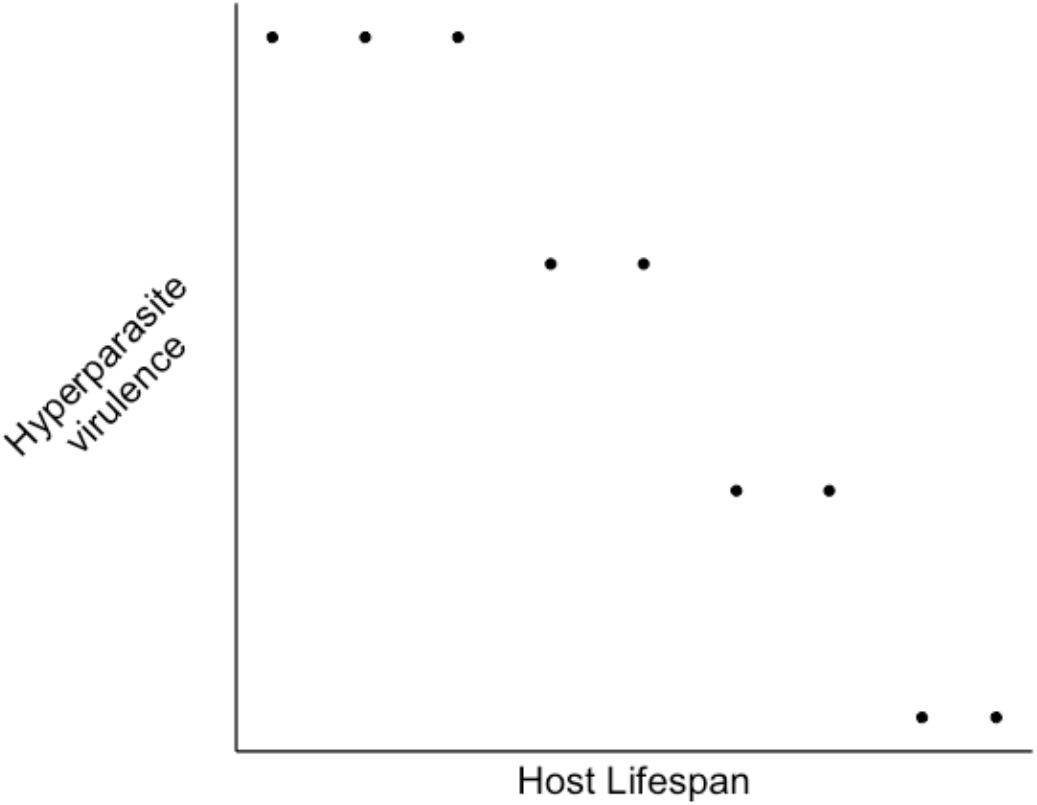
The relationship between the ESS value of α_H_, and different values of fixed model parameters. Here we show changes in hyperparasite virulence over changing host death rates (i.e. host lifespan). Non evolving parameter values are otherwise consistent with those used elsewhere.

## Discussion

We have presented a general model of hyperparasitic interactions that can be applied to a range of specific systems and gives us broad insights into the evolutionary dynamics of hyperparasites. The model makes explicit the unique evolutionary dynamics resulting from the interaction between two pathogens, one playing host to another, and sharing a base host. Understanding the evolution of the hyperparasite, and specifically the evolution of the effect of the hyperparasite on the parasite’s traits, is key to understanding these systems more generally. We are only just beginning to uncover the true diversity of hyperparasites in natural populations and to date, little is known about the ecological and evolutionary consequences of hyperparasitism in nature. Our model can guide empirical work by identifying key life-history traits and hypotheses to be tested.

We have shown that the virulence of hyperparasites (and accordingly, their effects on the parasite’s (their host) natural history parameters) are sensitive to changes in parasite and host parameters as well as the key characteristics of the hyperparasitic interaction. Importantly, the extent to which hyperparasites are transmitted with their parasites – hitchhiking – is critical to the evolutionary dynamics in the hyperparasite. Our key result is that increased proportions of hitchhiking, *p*, by a hyperparasite causes selection for lower levels of hyperparasite virulence (*α_H_*), all the way to when *p* = 1 (hitchhiking every time) when avirulence is selected for. This makes intuitive sense when considering the virulence transmission tradeoff implemented in the model. We expect both virulence and transmission to be functions of host exploitation, with lower virulence and higher transmission being favorable (hence the tradeoff, as virulence decreases, so does transmission). However, when a hyperparasite is able to hitchhike, this leads to a different transmission pathway for the hyperparasite that is not subject to the virulence transmission tradeoff and therefore ‘cost free’. Effectively as hitchhiking increases, the fitness of the parasite and the hyperparasite are more tightly linked with both invested in parasite transmission, as the hyperparasite breaks out its own classical virulence-transmission trade off. This may also have other impacts on the relationship between parasite and hyperparasite. In addition to lower virulence, in systems where hitchhiking occurs, the negative impact of the hyperparasite on its parasite host may take place during life-history stages that take place between transmission seasons (Tollenaere et al. 2014). Here we do not examine hitchhiking as an evolvable trait as we believe it is most likely a consequence of the nature of a particular system (i.e. type of parasite, hyperparasite, specific biology), but our models emphasize that it is absolutely the critical trait determining the evolution of hyperparasite systems. The proportion of hyperparasite infections that hitchhike at parasite transmission events should therefore be a key trait that is estimated in the field when studying a natural hyperparasite system.

We examined the evolution of the impact of the hyperparasite on its parasitic host assuming that there are tradeoffs between hyperparasite traits as a function of host exploitation similar to the classic assumption of the trade-off theory of parasite virulence and transmission (Anderson and May 1982, Alizon et al. 2009). The conceptual basis of these trade-offs is that the growth of the hyperparasite in the parasite causes harm that could reduce the growth rate of the parasite in the host and therefore reducing its virulence (decreasing ***c_α_*** to between 0 and 1) transmission (decreasing ***c_β_*** to between 0 and 1) and making it easier for the host to clear (increasing ***c_γ_*** above 1). This is the same conceptual basis to the classic trade-off assumption for which there is mounting empirical evidence (Acevedo et al. 2019, Atkins et al. 2011). In particular, we focused on values of *c_α_*, the parasites reduction in virulence as a result of hyperinfection: the induction of “hypovirulence”. The idea that hyperparasites can select for hypovirulence in their hosts has been the subject of considerable attention in specific systems (Choi and Nuss 1992, Milgroom and Cortesi 2004) but here we examine the general conditions under which it is likely to be found. Selection for hypovirulence increases in strength as parasite virulence on its host increases and as such, we expect the evolution of hypovirulence in hyperparasites of highly virulent parasites. Hypovirulence is also selected for when hyperparasites cannot hitchhike, as there is selection for higher levels of hyperparasite virulence and transmission in this case (and therefore also values of *c_α_* that are smaller). Parasites with acute infections (high recovery rates) also select for hypovirulence. Hyperparasitic biocontrol agents aimed at reducing the virulence of the parasites would therefore be most likely to be found for non-hitchhiking, highly virulent acute parasites. In natural communities, hypovirulence inducing hyperparasite could promote host-parasite coexistence by attenuating parasite virulence (Weldon et al. 2013).

The traits of the parasites are also important in selecting their hyperparasites. In particular, we show that in acute parasites, where the host is more able to clear infection, will select for decreased hyperparasite virulence. This makes sense because both of these processes take hyperinfected host individuals and send them back to the susceptible class. Furthermore, parasites with higher virulence select for hyperparasites of higher virulence as they prioritize transmission given the shorter lifetime (infectious period) of their parasite host. The interesting implication of this is that we expect high hyperparasite virulence when they infect highly virulent pathogens that the host finds difficult to clear. This is the result of the unique nature of hyper-parasite rather than parasitic infection and it points to the role that hyperparasites may play in combating such dangerous parasites.

We have also shown how host traits can also influence selection pressures on the hyperparasite. Despite not being the hyperparasite’s direct host, the parasite-hyperparasite interactions occur in the context of the base host. We show that if this host is particularly long lived this may result in selection for lower levels of virulence in the hyperparasite. It is therefore important to not only consider the parasite and hyperparasite traits as important for selection but to look at the tripartite system in its entirety. It follows that a knowledge of the host life history can improve our ability to make predictions of the nature of the parasite-hyperparasite interaction. These predictions may allow the identification of systems which may be susceptible to hyperparasitic invasion or candidates for biocontrol of a parasite (Rigling and Propsero 2017).

Although there is limited data, we are able to compare our predictions to what we see in hyperparasitic systems in nature. Table 2 contains estimated parameter values from the literature for several notable examples of hyperparasitic interactions. Based on these values, we can see that estimated relative magnitudes of *p, c_α_, c_β_, c_γ_* seem to mimic the predictions of our model at least qualitatively. For systems with observed low amounts of hitchhiking, there are as we predict comparatively higher levels of virulence. Conversely for systems where we expect hitchhiking *p* to be larger, the resulting levels of hyperparasite virulence do indeed seem to be lower in nature. It is important to note that we cannot say anything about precise values of these parameters as the exact tradeoff functions or parameter values are not well known. But our models emphasize that determining whether a hyperparasite is a hitchhiking hyperparasite allows us to make several strong predictions about what we should see about the other parameter values of interest.

Interest in *C. parasitica* and CHV-1 stems primarily from the observation that CHV-1 can reduce the virulence of *C. parasitica* in its host the chestnut tree (Choi and Nuss 1992). Our model would predict that this will occur in CHV-1 in a system where *p* is close to 1, and if *C. parasitica* is highly virulent. There has been previous work investigating how not all strains of CHV-1 may induce hypo-virulence in *C. parasitica* and can be the result of be a combination of biological factors (Milgroom and Cortesi 2004). Further it has been shown that in some cases, hypovirulence inducing strains of CHV-1 can be outcompeted by other strains (Bryner and Rigling 2012). It is outside the scope of the current model presented here to specifically address when CHV-1 causes a reduction in virulence, but models incorporating vegetative compatibility (i.e. the phenotypic diversity and population substructure) (Milgroom and Cortesi 1999) and more specific genetic assumptions could further illuminate this. While there is strong evidence to suggest there is variation amongst CHV-1 strains in transmission (Deng et al. 2009, Ding et al 2007), there have been no studies done on variation of hitchhiking ability between strains. If evidence could be found that there is variation between CHV-1 strains in hitchhiking, further modeling work may be needed to investigate how selection may occur on this trait. What is clear when examining the *C. parasitica* and CHV-1 system is that *C. parasitica* can be highly virulent in natural settings (Peever et al. 2000), and it has been shown that naturally occurring CHV-1 may have been a factor in differences in outcome at the population level (Bryner et al. 2012). As discussed previously our model shows that the high virulence of *C. parasitica* provides pressure to drive the system towards the strategy employed by CHV-1 in natural settings. The use of CHV-1 in targeted biocontrol strategies has the potential to lead to artificial selection but on traits beneficial for this use. This could provide a good test of our model. Overall, it is not surprising to find that the effect of hyperparasites is not consistent across space and time, as these interactions may be strongly mediated by the local environment (Zewdie et al 2021).

Other prominent examples of hyperparasitic systems suggest that the conclusions derived from this model hold in multiple systems (Table 2). For example, fungal hyperparasites, such as members of the *Ampelomyces* genus, tend to have low *p* and therefore select for higher values of *α_H_*. This is similar to lytic bacteriophage systems which we would also expect *p* to be low but observed virulence can be extremely high although given that they are obligate killers direct inference is difficult. Lysogenic phages in contrast are often transmitted with their bacterial hosts and we would therefore predict very different impacts of the lysogenic compared with lytic phage. Adapting our modelling framework to explore the range of phage-bacteria systems is likely to reveal further nuances of coevolutionary dynamics. There is a great diversity of hyperparasites in nature, but relatively little is known about the key parameters we have identified as important for understanding host-parasite-hyperparasite evolution. This paper provides motivation for studies to estimate more of these parameters in natural systems. In particular there is a critical need to measure the probability of hitchhiking, which our models have shown to have fundamental effect on resulting selection.

In summary, we have presented a general model of the evolution of hyperparasites using adaptive dynamics. We showed that the ability of the hyperparasite to hitchhike with a primary infection can have dramatic effects of the ES levels of virulence and transmission of the hyperparasite. We also show how the life history traits of the intermediate parasite can exert effects over selection on the hyperparasite showing how hyperparasite systems can have very different evolutionary behavior despite having a similar, tri-species, hierarchical structure. Ultimately our results can inform the conditions under which we might expect induction of hypovirulence by a hyperparasite, with implications for biocontrol.

## Acknowledgments

This work was completed with funding support from grants from the NIH R01 GM122061-03 and the NSF DEB-2011109 to MB.

## Data Accessibility Statement

Code to reproduce figures and numerical analysis is available at github.com/gnorthrup/HyperparasiteEvolution

