## Supplemental Information for "The Evolutionary Dynamics of Hyperparasites"

**The Evolutionary Dynamics of Hyperparasites Supplemental Information**

**Invasion Equations**

In order to determine which hyperparasite is more fit between a mutant and a wild type with differing parameter values, we must create a model which incorporates both. This is done by introducing a fourth equation to the system seen in the main text (1). By adding an $M$ class (hyperparasitized with mutant hyperparasite class) we get a system of equations shown below in (2).

$$\frac{dS}{dt} = (b(1-qN)-d)S - \beta_I SI - c_\beta \beta_I SH - c_{\beta M}\beta_I SM + \gamma_I I + (c_\gamma \gamma_I + \alpha_H)H + (c_{\gamma M}\gamma_I + \alpha_M)M$$

$$\frac{dI}{dt} = \beta_I SI + (1-p)c_\beta \beta_I SH + (1-p_M)c_{\beta M}\beta_I SM + \gamma_H H + \gamma_M M - \beta_H IH - \beta_M IM - (\alpha_I + \gamma_I + d)I$$

$$\frac{dH}{dt} = pc_\beta \beta_I SH + \beta_H IH - (c_\alpha \alpha_I + c_\gamma \gamma_I + d + \alpha_H + \gamma_H)H \tag{2}$$

$$\frac{dM}{dt} = p_M c_{\beta M}\beta_I SM + \beta_M IM - (c_{\alpha M}\alpha_I + c_{\gamma M}\gamma_I + d + \alpha_M + \gamma_M)M$$

Here, we also introduce a set of parameters with subscript M, indicating this is a parameter dealing with the mutant hyperparasite. This system can then be used to determine which hyperparasite is more fit, for a variety of parameter values. For simpler systems this can be done analytically; however, the high amount non-linearities of this system renders the analytical approach computationally intractable, so a numerical approach was better suited.

**Numerical Analysis of Fitness**

To determine the more fit hyperparasite for a given parameter combination, we analyzed the trajectory of the system by first allowing it to equilibrate in the absence of mutants. This allows us to verify the assumption of the system being at equilibrium at the introduction of the mutant. Then after an introduction of the mutant into the system, the trajectory can be examined to determine if the original equilibrium was returned to (i.e. the S,I,H equilibrium is numerically stable) or if a new equilibrium was reached featuring the mutant hyperparasite class dominating (i.e. the S,I,H equilibrium is numerically unstable). The first case indicates the wild type hyperparasite is more fit, the second case indicates the mutant hyperparasite is more fit. This procedure is numerically equivalent to the purely analytical approach due to how fitness functions are calculated but allows for analysis of this more complex system in a straightforward way. We then repeat this across combinations of hyperparasite virulence ranging from 0 to 1 and the associated parameters to identify the parameters values  which can always invade as the mutant and cannot be invaded as the wild type, an evolutionarily stable strategy. This strategy is then repeatedly for a wide variety of parameter sets, to see how changing parameter values ultimately changes the evolutionary outcomes of the system.

**Trade-offs**

We used two different trade-offs in this paper, the first was a simple, saturating, virulence transmission trade-off. This was the trade-off used to create the evolutionary outcomes shown in **Figure 2**. We then included the parameters $c_\alpha, c_\beta, c_\gamma$ in this trade-off, as these can be other ways that exploitation by the hyperparasite can potentially decrease parasite fitness. We used a simple linear function for these traits in the trade-off, with $c_\alpha$ & $c_\beta$ linearly decreasing equal increases in $\alpha$. Recall that these parameters multiple the rate, so decreasing them means that the hyperparasite is reducing virulence or transmission by a larger amount. Conversely, we increase $c_\gamma$ as alpha increase, indicating that the host recovery rate from the parasite is increasing as hyperparasite replication increases.

**Parameter Values**

As this model is not built on or fit to a particular system, the parameter values ranges are set wide enough to encompass a variety of different biological systems, only specifying the requirement for stability of the tripartite equilibrium. Table S1 contains the non-evolving parameter values used in figures 3-5, which are plotted over. The values of ES hyperparasite virulence $\alpha_H$ always range on 0 to 1.

| Parameter | Range of values |
|---|---|
| Parasite virulence, $\alpha_I$ | 0.00-0.20 |
| Hitchhiking probability, $p$ | 0-1 |
| Host recovery rate, $\gamma_I$ | 0-3 |
| Host lifespan, $d$ | 0.01-0.4 |

*Table S1: Non-evolving parameter ranges over which ESSs are calculated and plotted in figures 3,4 and 5*

.
